# Structure and lipid dynamics in the *A. baumannii* maintenance of lipid asymmetry (MLA) inner membrane complex

**DOI:** 10.1101/2020.05.30.125013

**Authors:** Daniel Mann, Junping Fan, Daniel P. Farrell, Kamolrat Somboon, Andrew Muenks, Svetomir B. Tzokov, Syma Khalid, Frank Dimaio, Samuel I. Miller, Julien R. C. Bergeron

**Affiliations:** Department of Molecular Biology and Biotechnology, The University of Sheffield, Sheffield, United Kingdom; Randall Division of Cell and Molecular Biophysics, King’s College London, London, UK; Department of Microbiology, The University of Washington, Seattle, USA; Department of Biochemistry, The University of Washington, Seattle, USA; Department of Chemistry, University of Southampton, Southampton, UK; Department of Genetics, The University of Washington, Seattle, USA; Ernst-Ruska-Centre 3, Forschungszentrum Jülich, Germany; Department of Pharmacology, The University of Washington, Seattle, USA

## Abstract

Multi-resistant bacteria are a major threat in modern medicine. The gram-negative coccobacillus *Acinetobacter baumannii* currently leads the WHO list of pathogens in critical need for new therapeutic development. The maintenance of lipid asymmetry (MLA) protein complex is one of the core machineries that transport lipids from/to the outer membrane in gram-negative bacteria. It also contributes to broad-range antibiotic resistance in several pathogens, most prominently in *A. baumannii*. Nonetheless, the molecular details of its role in lipid transport has remained largely elusive.

Here, we report the cryo-EM structures of the core MLA complex, MlaBDEF, from the pathogen *A. baumannii*, in the apo-, ATP- and ADB-bound states. These structures reveal multiple lipid binding sites, in the cytosolic and periplasmic side of the complex. Molecular dynamics simulations suggest their potential trajectory across the membrane. Collectively with the recently-reported structures of the *E.coli* orthologue, these data also allows us to propose a molecular mechanism of lipid transport by the MLA system.

## Introduction

Gram-negative bacteria are enveloped by two lipid bilayers, separated by the periplasmic space containing the peptidoglycan cell wall. The two membranes have distinct lipid compositions: The inner membrane (IM) consists of glycerophospholipids, with both leaflets having similar compositions, while the outer membrane (OM) is asymmetric, with an outer leaflet of lipopolysaccharides (LPS) and an inner leaflet of glycerophospholipids (1) (Fig. 1 A). This lipid gradient, depicting the first and most important permeation barrier is maintained by several machineries, including YebT, PqiB, and the multicomponent MLA system (2, 3), which consists of MlaA present in the OM, the shuttle MlaC in the periplasmic space, and the MlaBDEF ABC transporter system in the IM (Fig. 1 A). The structure of some of these components have previously been solved: the OM protein MlaA, which was found to form a stable complex with outer membrane porins OmpF and OmpC (4, 5), and the periplasmic protein MlaC revealing a hydrophobic pocket for direct lipid transport through the periplasm (2, 6). Low-resolution cryo-EM maps of the MlaBDEF core complex, from *Escherichia coli* (MlaBDEF_ec_)(2) and *Acinetobacter baumannii* (MlaBDEF_ab_) (7) have also been reported, and revealed the overall architecture of the complex, but did not allow to elucidate the molecular details of lipid binding and transport. Opinions about the directionality of lipid transport by the MLA system have been highly controversial, with initial reports suggesting that it recycles lipids from the OM to the IM (2, 8, 9), but recent results (7, 10, 11) indicated that it might exports glycerophospholipids to the outer membrane.

**Figure 1:**
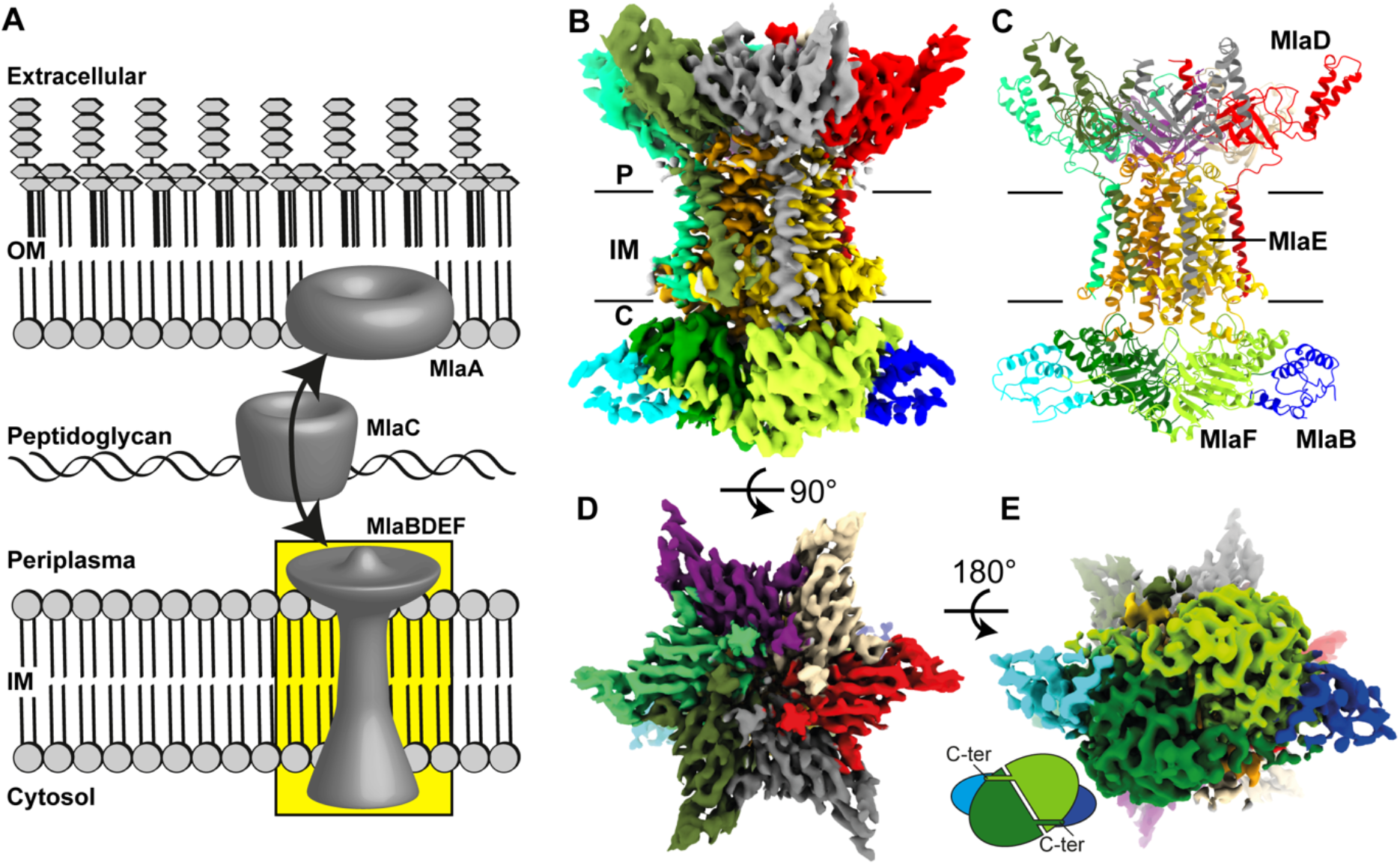
Structure of MlaBDEF_ab_. A: Scheme of lipid transport by the MlaABCDEF system in gram-negative bacteria (OM = outer membrane; IM = inner membrane). B: 3.9 Å CryoEM map of MlaBDEF_ab_-AppNHp in DDM (C=cytosol, IM=inner membrane, P=periplasma). C: MlaBDEF_ab_ has global C2 symmetry with six copies of MlaD that span the inner membrane from the periplasmic space to the cytosol, two copies of MlaE embedded in the membrane (yellow/gold), two copies of the ATPase MlaF in the cytosol (green), each bound to a copy of MlaB (blue/cyan). D: top view of the MlaBDEF_ab_ complex reveals C6 symmetry of MlaD. E: C-terminal regions of MlaF bind the opposing MlaB subunit via a “handshake-mechanism”.

In this study, we report the structure of the MlaBDEF_ab_ complex in detergent, in three nucleotide states, by single-particle cryo-EM. We also performed molecular dynamics simulations to gain insights into the dynamics of lipids within their observed binding sites. Collectively, this provides important new insights in the mechanism of lipid transport by the MLA system, and about the characterization of membrane proteins in detergent.

## Results

### Structure of MlaBDEF_ab_

We had previously reported the purification of MlaBDEF_ab_ in the presence of the detergent n-dodecyl β-D-maltoside (DDM), and its structure to ~ 8 Å, by single-particle cryo-EM, from data collected on a side-entry 200kV microscope (7). In order to improve the resolution of this structure, we collected a dataset of the same complex, in the presence of the non-hydrolizable ATP analogue App-NHp, using a state-of-the-art Titan Krios instrument. Using this better and larger dataset, we were able to refine the structure to ~ 3.9 Å resolution (Fig. 1B, Suppl. Fig. 1, Table 1). This map allowed us to build a *de-novo* atomic model using Rosetta (12) (See Materials and Methods for details).

**Table 1:** Cryo-EM data processing and refinement statistics.

|  | **AppNHp**<br>(EMDB-11082<br>(PDB: 6Z5U) | **Apo**<br>(EMDB-11083) | **ADP**<br>(EMDB-11084) |
| --- | --- | --- | --- |
| **Data Collection and Processing** |  |  |  |
| Microscope | Titan Krios | Titan Krios | Titan Krios |
| Voltage (kV) | 300 | 300 | 300 |
| Camera | K2 summit | K3 bioquantum | K2 summit |
| Pixel size (Å) | 1.07 | 0.41 | 1.07 |
| Defocus range (μm) | -1 to -2.5 | -0.8 to -2.0 | -1 to -2.5 |
| Total dose (eÅ<sup>-2</sup>) | 47 | 40 | 40 |
| Number of micrographs | 2557 | 4737 | 1901 |
| Total particles used | 93,295 | 28,699 | 62,512 |
| Map Resolution (Å) | 3.92 | 4.24 | 4.43 |
| **Refinement** |  |  |  |
| Model composition |  |  |  |
| Non-hydrogen atoms | 17,892 |  |  |
| Protein Residues | 2334 |  |  |
| Ligand atoms | 4 |  |  |
| *B* factors (Å<sup>-1</sup>) |  |  |  |
| Protein | 145.24 |  |  |
| Ligand | 170.31 |  |  |
| R.m.s. deviations |  |  |  |
| Bond lengths (Å) | 0.008 |  |  |
| Bond angles (°) | 1.384 |  |  |
| Validation |  |  |  |
| MolProbity score | 3.43 |  |  |
| Clashscore | 35.15 |  |  |
| Poor rotamers (%) | 10.94 |  |  |
| Ramachandran plot |  |  |  |
| Favoured (%) | 87.45 |  |  |
| Allowed (%) | 10.94 |  |  |
| Disallowed (%) | 0.43 |  |  |

As shown of figures 1B, and 1C, the transmembrane multiprotein complex features a 6-fold symmetric assembly of MlaD, with the C-terminal helix forming a basket in the periplasmic space (Fig. 1D), and the N-terminal helix spanning the inner membrane (Fig. 1C). The N-terminal TM helices of MlaD are wrapped around the two MlaE molecules in the membrane, with three MlaD helices interacting asymmetrically with one MlaE, as reported previously (7). Intriguingly, while MlaE was predicted to have 6 TM helices, we observe that TM1 does not traverse the membrane, but is monotropically embedded in the inner leaflet, a feature similar to the G5G8 human sterol exporter (13), suggesting similar mechanisms between these complexes, and further supporting the MLA system as a lipid exporter. On the cytosolic side, MlaE is anchored into the ATPase MlaF via the coupling helix situated between TM3 and TM4 (Fig. 1), again similar to the G5G8 sterol exporter complex. MlaF is bound to MlaB away from the nucleotide binding site, similar to the recently-reported *E.coli* MlaBF structure (14), with C-termini of MlaF binding the opposing MlaB subunit by a “handshake” mode (Fig. 1E). We note, however, that particle classification demonstrated that only ~50% of the particles included MlaB bound to both MlaF, leaving the other 50% bound to only one copy of MlaB (Suppl. Fig. 2). Dual binding of MlaB did not introduce major structural alterations at the detected resolution, and this observation may correspond to a regulatory role for MlaB or could be due to complex disassembly during sample preparation.

### Lipids spontaneously bind into pockets of cytoplasmic MlaBDEF_ab_

In the cryo-EM derived map of MlaBDEF_ab_, observed well-defined density in the pocket formed between the MlaE TM1, and two MlaD helices, within the inner leaflet of the lipid bilayer (Fig 2 A-C), which could not be interpreted by protein atoms. We also observed that this pocket is coated by cationic residues, mainly Arg14, Arg47 and Arg234 of MlaE, forming a charged pocket that might attract a lipid head group (Fig. 2D). This observation prompted us to propose that this density corresponds to detergent molecules bound to the complex. As the sample was solubilized in DDM, this density is most likely occupied by the maltoside ring, with the flexible hydrophobic chains extending up into an apolar region of MlaE. In order to confirm if the observed density was consistent with lipid positioning, we performed molecular dynamics simulations with unoccupied binding pockets as starting structures and observed rapid incorporation of bulk lipids during equilibration (Fig. 2 E). Newly bound lipids remained stably bound during 500 ns production runs (Fig. 2 F), depicting a reasonable first step in lipid export. This result is consistent with the position of lipid molecules within this region of the MlaBDEF_ab_ complex.

**Figure 2:**
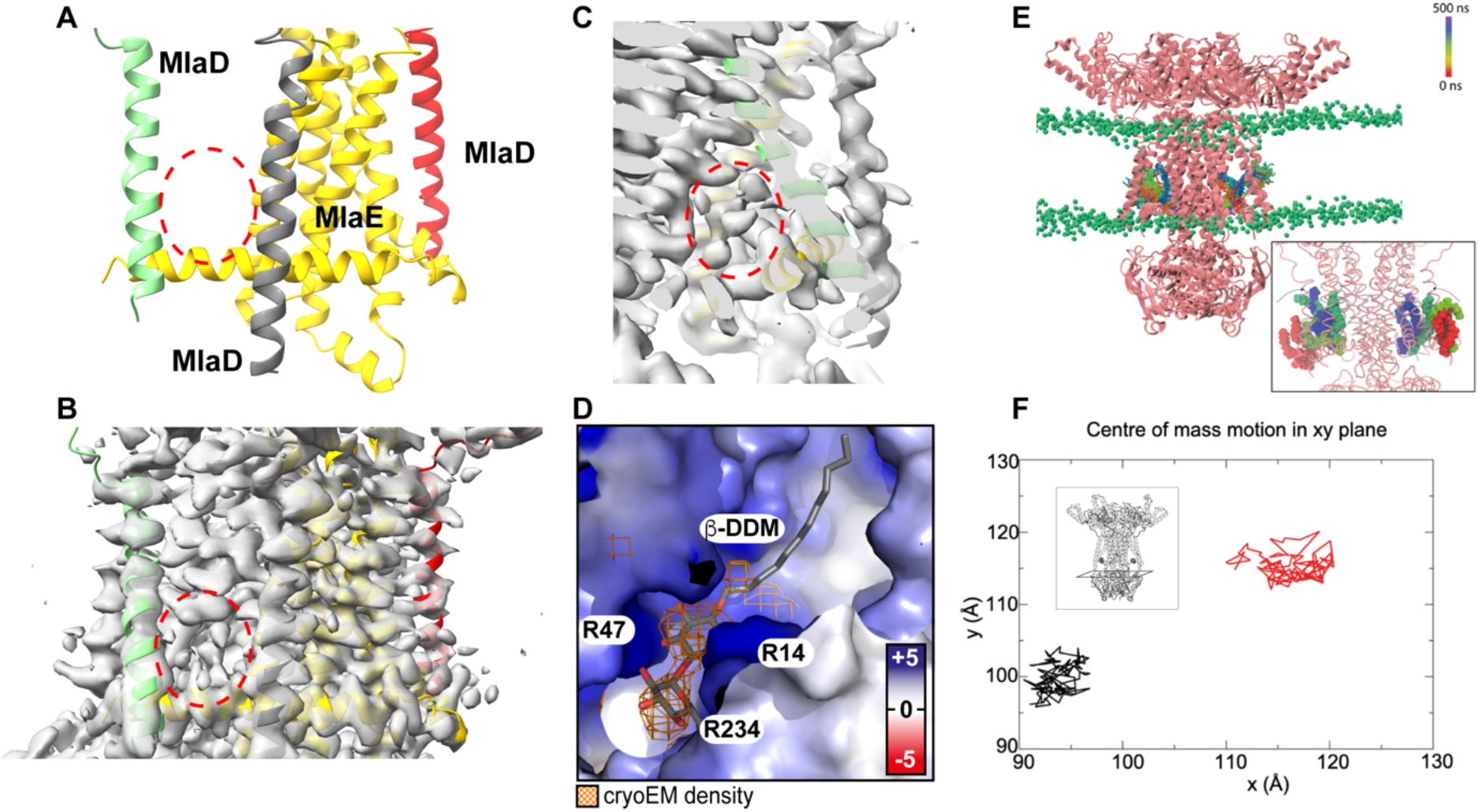
Detergent position and dynamics at the MlaD TM-MlaE interface. A-C: Transmembrane region of MlaBDEF_ab_ with one MlaE and three MlaD proteins shown. Red circles indicates density unoccupied by protein residues. D: Electrostatic surface potential of modeled inner membrane MlaBDEF_ab_ (blue=+5 kT/e, white=0kT/e, red=−5kT/e) shows a cationic binding site for anionic lipid head groups like □-DDM and an apolar pocket for the uncharged lipid tails. □-DDM was modeled inside the previously empty CryoEM density (orange). E-F: Molecular dynamics simulations show spontaneous, stable binding of lipids into the cytosolic binding pockets of MlaBDEF. Lipid-free MlaBDEF_ab_ was embedded into an inner membrane model system of 75% POPE, 20% POPG, 5% Cardiolipin (Methods section for details). Panel E shows the location of two POPE lipids during a 500 ns simulation, the colour scheme indicates the movement of the lipids as shown in the legend. In this simulation the lipids moved into this location spontaneously during the equilibration process, as shown in the close-up view in the inset in which the red coloured lipids indicate starting positions before the equilibration process. Panel F shows the motion of the lipids in the xy plane (inset) over 500 ns. The lipids are confined within an area of 10 x 10 Å for 500 ns, showing this location is favourable.

### Lipids bind into the periplasmic basket and partially flip

The periplasmic region of MlaBDEF_ab_ consists mainly of hexameric MlaD forming a basket shape (Fig. 1). Similar to other MCE domain proteins (15), MlaD consists of a central beta sheet motif with a central pore loop that is formed by hydrophobic Leu153/Leu154 in the center of the C6 symmetric complex (Fig. 3 A). We observed that in our map, unattributed density was present between the central pore loops, as well as in the central pore (Fig. 3 A). Importantly, this density is not an artifact of the 6-fold symmetry, as it is also resolved when no symmetry was applied during the reconstruction. The presence of lipid molecules in very similar positions were previously reported in the MCE protein YebT (16), which prompted us to postulate that this unattributed density on the MlaD periplasmic region also corresponds to detergent molecules. Of note, the basket region has previously been proposed to form the binding site for the periplasmic carrier protein MlaC (4), suggesting that lipids are extracted from this position upon MlaC binding, consistent with the interpretation that these regions of the map correspond to lipid molecules.

**Figure 3:**
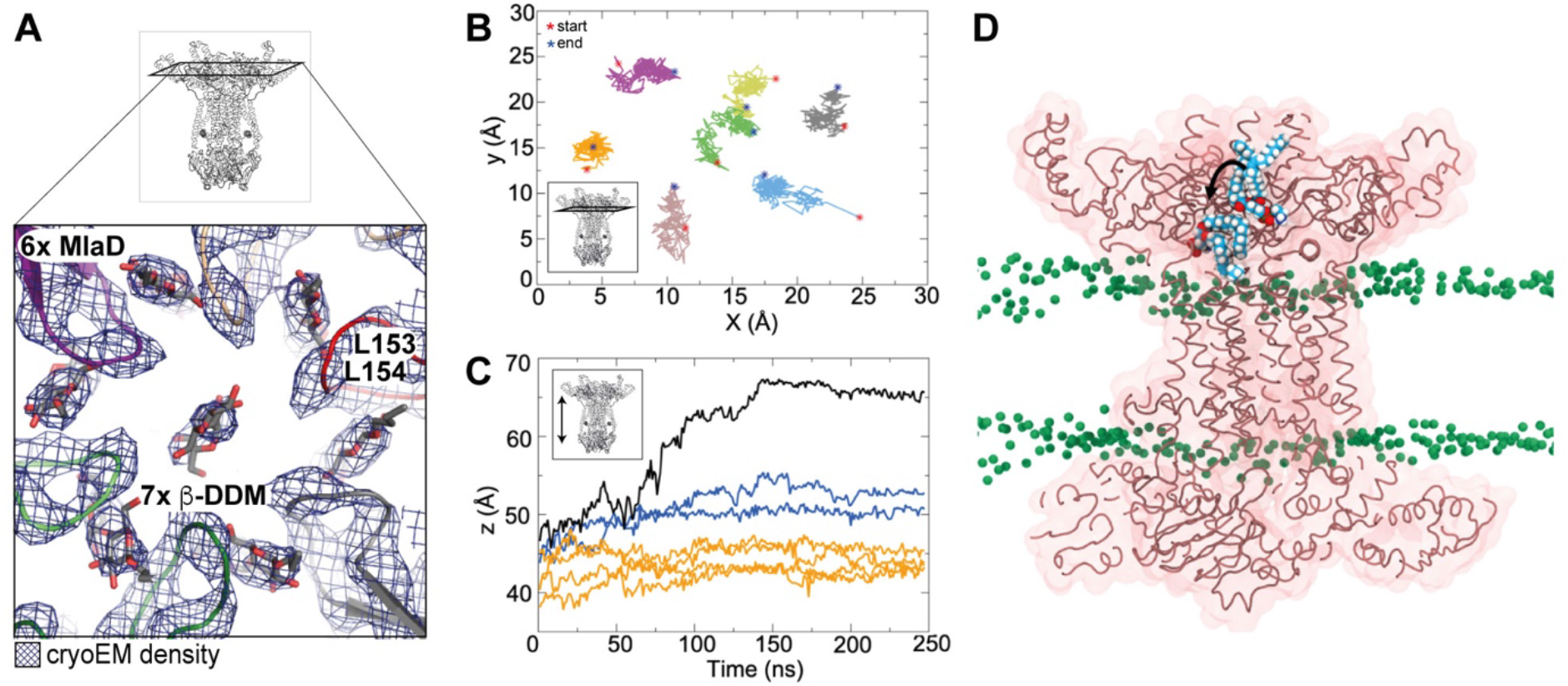
Lipid position and dynamics in the MlaD basket. A: Close-up of the periplasmic MlaD basket that houses six peripheral and one central detergent molecule. B-D: Spontaneous lipid flipping within the basket region of MlaBDEF_ab_ during 250 ns molecular dynamics simulations. Panel B shows the center of mass motion of the seven lipids in the xy plane. They are confined to an area of ~5 x 5 Å, indicating this is a high lipid affinity region. Panel C shows the center of mass movement of the seven lipids in the Z dimension as a function of time. The higher values of z correspond to the cytoplasmic end of the protein. The lipid shown in panel A corresponds to the black curve showing a clear movement towards the cytoplasmic end. Three other lipids (blue) move into this channel but to a lesser extent than the aforementioned lipid, whereas three others (orange) remain close to their starting positions. Simulations from a model at the reported resolution cannot clarify the directionality of the movement, but rather that these regions are conduits for lipids. Panel D shows a cut-away view of the protein with a POPE lipid at two time points during the simulation, time = 0 ns and 150 ns. The simulation was initiated with seven POPE lipids placed at the periplasmic end of the protein corresponding to the density for detergents in the cryo-EM data (Table S2). The lipid which is displaced more towards the cytoplasmic end is from the frame at t = 150 ns. The central hydrophobic ‘channel’ of the protein is a clear conduit for lipids given the spontaneous movement of POPE into this region in just 150 ns.

Because of this likely important role of lipid dynamics for this region, we next performed 3D variability analysis (17), to identify molecular motions in the MlaD crown. As shown on Suppl. movies 1 and 2, this analysis revealed that the 6x MlaD crown can be translated and rotated against the MlaBEF transmembrane part, which may play a role in the lipid transport mechanism. Incidentally, this dynamic property likely also limits the achievable resolution for this region of the map.

To further investigate the dynamic properties of the lipids present in the MlaD basket, we subjected them to molecular dynamics simulations. As shown on Figures 3 B-C, we observed that the peripheral lipids are rather stable during the course of the simulation. In contrast, the central lipid undergoes a significant motion within 150 ns of the trajectory. Remarkably, during this motion, we observed lipid flipping (Figure 3 D), with the head group which was modeled away from the central pore operating an almost 180°, with the polar group buried within the MlaE transporter. This observation likely indicates that we had initially build the lipid in the wrong orientation, and highlights the importance of properly modeling lipid molecules, especially at the intermediate resolution such as that of our MlaBDEF_ab_ map. In addition, this demonstrates the presence of a large hydrophobic pocket at the MlaD-MlaE interface, which could correspond to the channel for lipid transport.

### Nucleotide binding occurs at the interface of MlaE, MlaF and MlaD

As indicated above, our structure of MlaBDEF_ab_ was determined in the presence of the non-hydrolizable ATP analogue AppNHp. Accordingly, we observed clear density of the nucleotide and Magnesium, within the MlaF binding pocket (Fig. 4A). The nucleotide links the ATPase subunits with the lipid transport domains at the interface of MlaF, MlaE and the N-terminus of MlaD. The resolution was sufficient to map the phosphate binding regions like Walker-A motif around Lys55 of MlaF, as well as binding of the *γ*-Phosphate by Ser51 and His211 (H-loop). The Mg^2+^-ion is coordinated by the Walker-B motif around Asp177 (Fig. 4A) and adenosine ring binding is achieved through Arg26 in the A-loop. As revealed in the previously-reported low-resolution map (7), the position of MlaF suggest that the complex is in the substrate-bound conformation.

**Figure 4:**
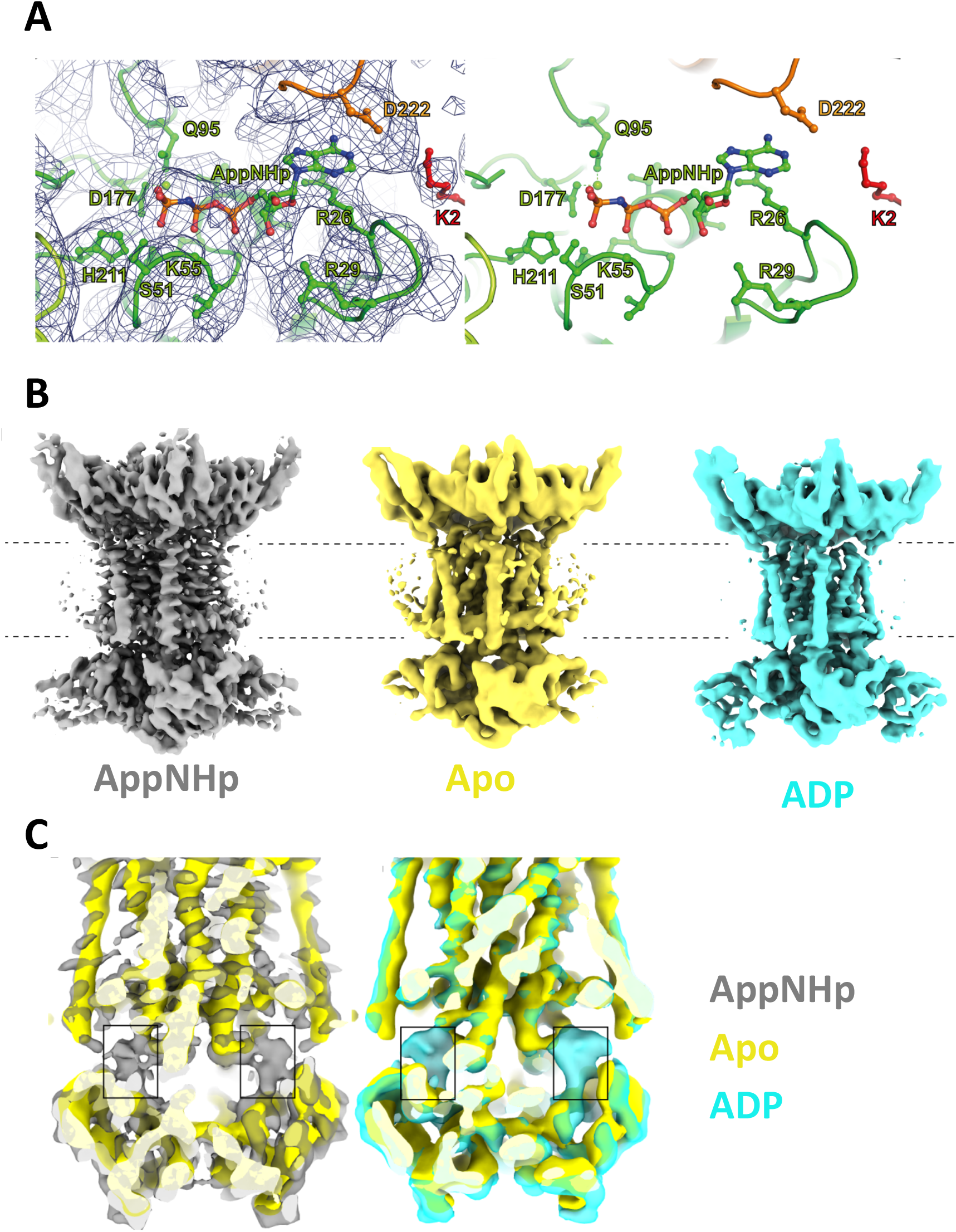
Nucleotide position in the MlaBDEF_ab_ structure. (A) AppNHp is bound at the interface of MlaE (orange), MlaD (red) and MlaF (green). B: maps of MlaBDEF_ab_ bound to AppNHp (grey), its *apo* state (yellow) and bound to ADP (cyan). (C) The overlay of the maps unambiguously confirms the presence of nucleotide in the AppNHp and ADP maps, but not in the Apo map. Nonetheless, no overall structural changes is observed.

In order to identify structural changes in the complex associated with ATP binding and hydrolysis, we next determined the structure of MlaBDEF_ab_ without nucleotide (Suppl. Fig. 3), and with ADP (Suppl. Fig. 4). As shown on Figure 4A-C, we obtained both structures, to ~4.2 Å and ~ 4.4 A, respectively (Table 1). We note that in spite of the nominar global resolution, the map for the complex bound to ADP possesses features largely similar to that of the complex bound to AppNHp. In contrast, the map of the apo complex is less well resolved, in particular with most TM helices being mostly featureless. This suggests that nucleotide binding stabilizes the overall architecture of the MlaBDEF_ab_ complex.

Nonetheless, we note that the overlay of the MlaBDEF_ab_ maps in the apo, AppNHp-bound, and ADP-bound states show no major structural changes in any of the complex components, other than the nucleotide binding site (Fig. 4D). This indicates that in spite of the nucleotides being bound to the ATPase domain, under the conditions used this is not sufficient to trigger the activation of the channel opening.

## Discussion

While this manuscript was under review, three independent groups released the structure of the *Escherichia coli* MlaBDEF complex (MlaBDEF_ec_), in a range of nucleotide states and with various solubilization approaches (18–20).

Comparison of the MlaBDEF_ec_ structure (PDB-ID 7CH0, 7CGN), determined in lipid nanodiscs, and the MlaBDEF_ab_ structure (this study), determined in DDM, reveals a ~10 Ang movement of the MlaD TM helices, accompanied by a ~30° tilt of the MlaE N-terminal helix (Fig. 5 A). This movement narrows the cytoplasmic allocrit binding pocket significantly (Fig. 5 A, red circles). A structural flexibility like this would allow adjustment of MlaBDEF for allocrits with varying sizes. However, as lateral pressure in the membrane plane of a detergent micelle is not comparable to a lipid nanodisc, differences in sample preparation can also cause movement of this hinge. Furthermore, although the MLA system is conserved in gram-negative bacteria, the sequence identity between the MlaBDEF_ab_ MlaBDEF_ec_ proteins is around 30-40% identity (depending on the protein), and it is therefore possible that the differences observed between the two structures correspond to variability between bacterial species.

**Figure 5:**
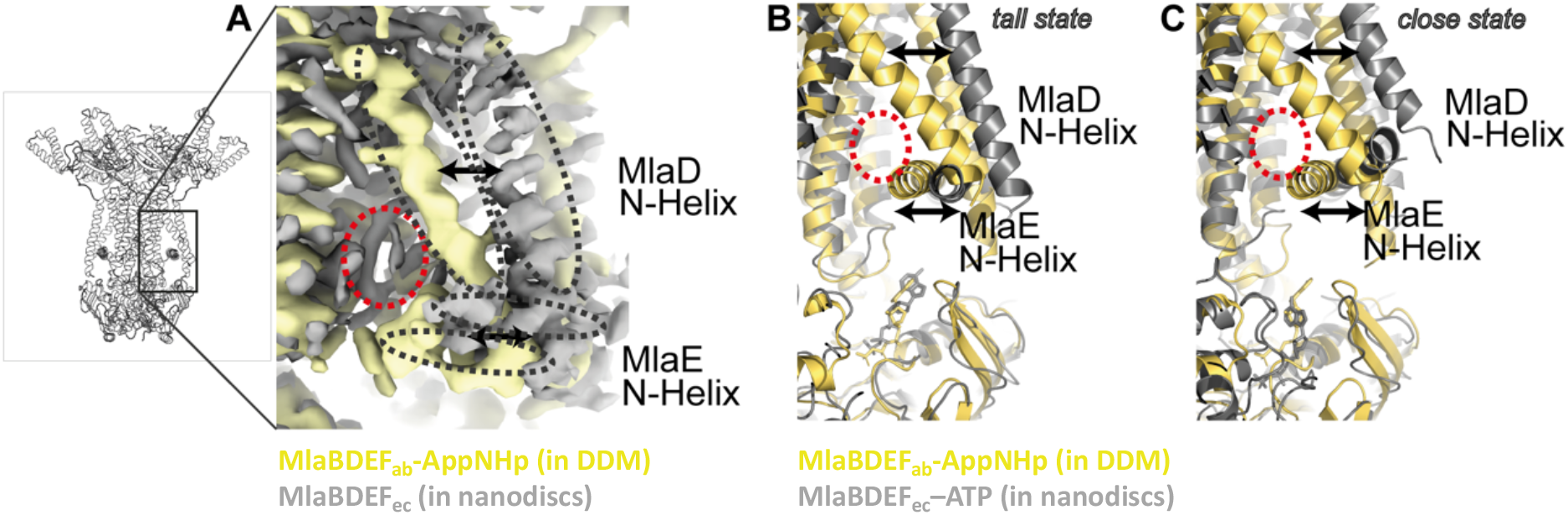
Comparison of the MlaBDEF_ab_ and MlaBDEF_ec_ structures. Acquired Cryo-EM map of MlaBDEF_ec_ in lipid nanodiscs (20) and MlaBDEF_ab_ in a DDM micelle (this study) show large differences in the lower lipid binding sites. A: Both the MlaD N-terminal helix and the MlaE N-terminal helix form a much smaller lipid binding pocket (red circle) in detergent environment (yellow, this study) compared to lipid environment (grey, EMD-30355). Two nucleotide binding states were observed in the MlaBDEF_ec_ study; “tall” and “close”. Comparison to B) “tall state” (grey) shows almost identical nucleotide position between MlaBDEF_ec_ and MlaBDEF_ab_, whereas

Significantly, in one of the aforementioned studies, two ATP-bound conformational states were observed for MlaBDEF_ec_: a more open conformation (“*tall state*”) and a tightly bound conformation (“*close state*”), with a significantly shifted nucleotide position. The nucleotide position in MlaBDEF_ab_ in detergent resembles the *tall state* with an open conformation of MlaF (Fig. 5 B-C).

We also note that the lipid binding pockets observed in our MlaBDEF_ab_ structures are also confirmed in the MlaBDEF_ec_ structures, notably the lipids found between the MlaD TM and the N-terminal helix of MlaE (20), and the lipid present in the central cavity (18–20). For the later, different exact lipid positions are observed depending on the state and study, but they are largely consistent with the lipid position obtained in the MD simulation, with the charged group buried at the MlaE-MlaD interface. In contrast, our map is the only one with clear lipid density at the interface between MlaD molecules in the periplasmic site; this could correspond to species specificity, or could be an artifact of the use of detergent for structure determination.

In the light of the multiple structures now available, the structures of MlaBDEF_ab_ reported here support a mechanism for lipid export as summarized in Figure 6. We observe lipids binding to the inner lipid binding pocket; this region is likely flexible and can possibly adapt to the size of the allocrit. MlaE is in very close proximity to ATP in the closed conformation, possibly describing an additional regulatory mechanism in *A. baumannii*. Structures of MlaBDEF_ec_ also showed a large-scale structural alteration upon tight ATP binding (20) that might be restrained due to the detergent environment in our experiments. We could furthermore observe a central lipid binding cite in the periplasmic basket region of MlaD that could partially flip into the central channel of MlaE during the course of our MD simulation. Nonetheless, neither our structures of MlaBDEF_ab_ nor the aforementioned MlaBDEF_ec_ could resolve allocrit transition between the cytoplasmic lipid binding pocket and the central channel. Similarly, allocrit transport from MlaBDEF to MlaC remains elusive.

**Figure 6:**
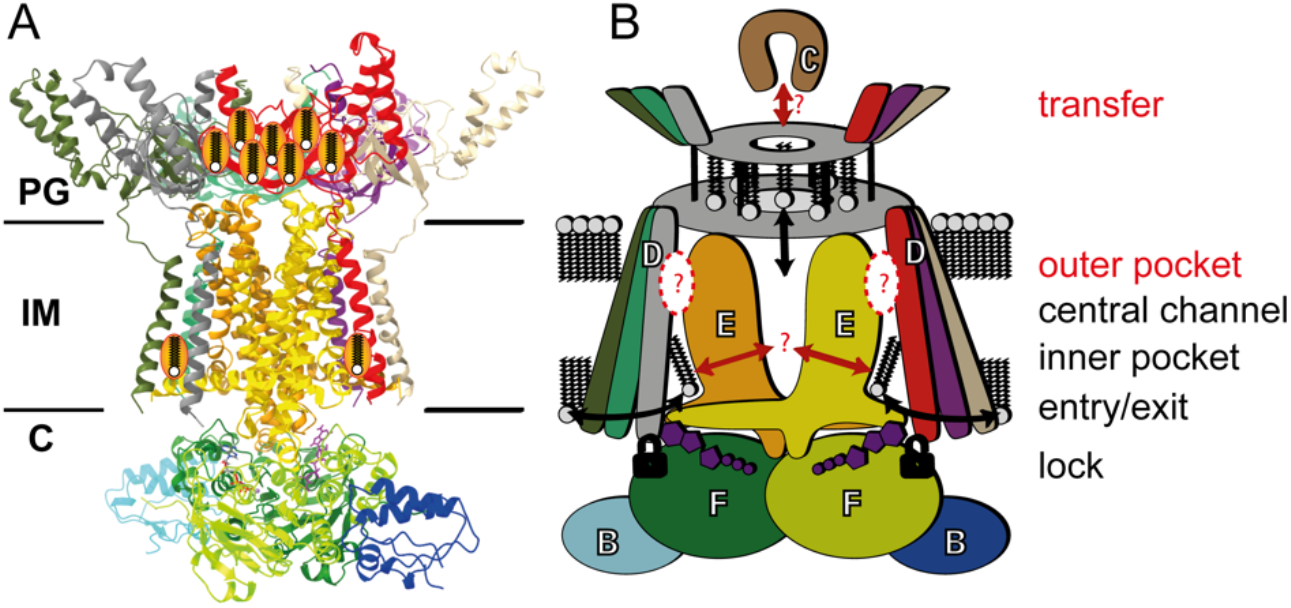
Molecular model of lipid transport by the MlaBDEF complex. A: structural model and B: observed degrees of freedom in this study. Lipids can move freely into the cytosolic paired binding pockets (“entry/exit”) that are very close to the ATP binding sites that directly bind MlaD (“lock”). Lipids can move and even flip inside the basket region (“transport”). C: The elucidated structures could not show how transport from the entry sites to the basket region is performed and how lipid exchange to/from MlaC is achieved.

It should be emphasized that the structural studies reported here support an anteretrograde direction for lipid transport, as supported by our previous biochemical data in *A. baumannii* (19) as well as other studies in *E. coli* (11, 19, 20). Nonetheless, other assays point towards a retrograde directionality (18, 21), and the current structures do not conclusively preclude this. Further structural analyses, in particular in complex with the periplasmic carrier MlaC, will be required to definitely resolve this controversy.

## Materials & Methods

### MlaBDEF_ab_ protein expression and purification

The expression and purification of MlaBDEF_ab_ has been described previously (7). Briefly, the plasmid was transformed into *Escherichia coli* BL21 DE3 cells and grown at 37°C until the cell density reached OD(600nm) = 1.0. The temperature was reduced to 20°C before induction with 1 mM isopropyl β-D-thiogalactoside (IPTG) and incubation overnight. Cells were harvested using centrifugation at 5000 x g and resuspended in ice-cold buffer A (20 mM Tris-HCl (pH 8.0), 150 mM NaCl, 5% (v/v) glycerol) before disruption in an ultrasonicator on ice (6 cycles; 60s run, 30s cooldown). Cell debris was centrifuged down at 17,000 x g for 10 min and the supernatant was centrifuged at 100,000 x g for 1h. This pellet was resuspended by gentle stirring in Buffer A supplemented with 1% (w/v) dodecyl-β-d-maltopyranoside (DDM) at 7°C for 1 h. After another centrifugation step at 100,000 x g for 30 min the supernatant was applied to a 5 ml Ni-NTA superflow column (GE Healthsciences) that was previously equilibrated with ddH_2_O and buffer A supplemented with 20 mM imidazole and 0.025% (w/v) DDM. The column was washed with buffer A supplemented with 20 mM imidazole and 0.025% (w/v) DDM before elution with buffer A supplemented with 300 mM imidazole and 0.025 % (w/v) DDM was performed. The elute was concentrated to 5 ml and applied to a gel filtration run on a 16-600 HiLoad Superdex 200 pg column (GE Healthcare), preequilibrated with 20 mM Hepes (pH 7.0), 150 mM NaCl, 0.025% (w/v) DDM. Peak fractions were collected, purest fractions were selected using SDS-PAGE and concentrated to 5 mg/ml.

### CryoEM sample preparation, data acquisition and image processing

4 μl freshly purified 5 mg/ml MlaBDEF_ab_ in 20 mM Hepes (pH 7.0), 150 mM NaCl, 0.025% (w/v) DDM was applied on 30s freshly glow discharged 300 mesh Quantifoil R2/2 grids, blotted for 3.5 s in a Leica EM-GP plunge freezer at 80% humidity and 4°C, before getting plunged into liquid ethane at −170 °C. For investigation of the AppNHp and ADP states, the protein was mixed 1:1 with 20 mM of the corresponding nucleotide in 50 mM Hepes (pH 8.0), 150 mM NaCl, 0.025 % (w/v) DDM buffer and incubated for 60 min on ice prior to grid preparation.

Micrographs of MlaBDEF_ab_-AppNHp were recorded on a 300 kV Titan Krios microscope with a Gatan K2 Summit detector in counting mode. 2557 movies were recorded with a pixel size of 1.07 Ang in 47 frames with 1e^−^/Ang^2^/frame. 1901 movies of MlaBDEF_ab_-ADP were recorded on the same instrument with a total dose of 40 e^−^/Ang^2^. The MlaBDEF_ab_ apo dataset was recorded on a 300 KV Titan Krios equipped with a Gatan K3 bioquantum detector. 4737 micrographs with a pixel size of 0.41 Ang were recorded with a total dose of 40 e^−^/Ang^2^ in 50 frames. Data processing was performed in CryoSPARC v2.14.2 (see Suppl. Fig. 1 and 3 for details). Raw images of the MlaBDEF_ab_-AppNHp dataset are publicly available under EMPIAR-10425. Sharpened maps in C2, C1 and C6 symmetry, masks and half-maps of MlaBDEF_ab_-AppNHp, ADP and apo are publicly available under EMD-11082, EMD-11083 and EMD-11084, respectively.

### Model building

The structural model of MlaBDEF_ab_ was manually built into the high-resolved regions of the MlaBDEF_ab_-AppNHp map using Coot. Computational intervention was required to build residues 4-38, and 94-134 of MlaD, as well as refine the AppNHp binding site of MlaF. RosettaES (12) was used to model residues 4-38 for each of the 6 MlaD subunits using the manually built model as the starting point. Residues 94-134 were first modeled with Rosetta *Ab-initio* (22) which yielded a tightly converged ensemble with a 2-helix topology. The top scoring model from the *Ab-initio* predictions was unambiguously docked into the corresponding density for each of the MlaD subunits using UCSF Chimera (23). Loops were completed and the entire MlaD crown refined using RosettaCM (24) with the context of the cryoEM density. To refine the AppNHp binding site of MlaF we first used RosettaCM to hybridize the manually built model with the homologous structures of (3fvq, chain A), and (4ki0, chain B) in the cryoEM density. Then using homology models and the unexplained density as a guide, a modified version of the rosetta protocol GALigandDock (unpublished) that uses the cryoEM data to drive sampling was used to dock the AppNHp molecule. As input to the protocol, a mol2 file of AppNHp was modified using openbabel (v. 2.4.1) (25) to add hydrogens, charges were assigned with the AM1-BCC charge method in antechamber (26) and a params file was generated with the script main/source/scripts/python/public/generic_potential/mol2genparams.py which is distributed with Rosetta. Finally, the entire complex was refined using RosettaCM, and Magnesium atoms were added by incorporating distance and angle constraints between the Mg atoms and the AppNHp oxygens on the β and γ phosphates during the final minimization. ISOLDE in ChimeraX was used to manually correct for modeling errors. Phenix.real.space.refine was used for refinement and validation. As shown on Suppl. Fig. 5 and 6, the resoluting model has an excellent fit to the map density, with clear fit of the side-chains in particular for the TM helices of MlaE and MlaD, which were critical for determining their registry. We note nonetheless that the TM for one of the MlaD molecules is mostly featureless (orange in suppl. Fig. 6), in which case we relied on the other two copies to position the helix in the density. The MlaBDEF_ab_-AppNHp model is publicly available under PDB-6Z5U.

### Molecular dynamics system preparation

The MlaBDEF_ab_ structure was completed by adding the missing residues using Modeller 9.23 (http://salilab.org/modeller/)(27, 28). A system, having the completed protein structure embedded in an inner *E.coli* innermembrane, was generated with the CHARMM GUI web interface(29–35) to have dimension of approximately 21 × 21 × 18 nm in *xyz* dimension. The innermembrane composition was based on previous atomistic simulations studies(36): 75% 1-palmitoyl-2-oleoyl-sn-glycero-3-phosphoethanolamine (POPE), 20% 1-palmitoyl-2-oleoyl-sn-glycero-3-phosphoglycerol (POPG) and 5% 1′,3′-bis[1,2-dioleoyl-sn-glycerol-3-phospho-]-sn-glycerol (Cardiolipin). To relax any steric conflicts within the system generated during set up, energy minimization of 5000 steps was performed on the starting conformation using the steepest descent method. An equilibration procedure followed in which the protein was subjected to position restraints with different force constant. The full equilibration protocol is shown in Suppl. Table 1. Fatty acid molecules were placed around the entrance of MLA using Visual Molecular Dynamics (VMD)(37).

### Equilibrium molecular dynamics simulations

All simulations were carried out with GROMACS 2019.6(38) version (www.gromacs.org) and CHARMM36(39) forcefield. For the nonbonded interactions and the short-range electrostatics cut offs of 1.2 nm was applied to the system with the potential shift Verlet cut off scheme, whereas the long-range electrostatic interactions were treated using the Particle Mesh Ewald (PME) method(40). All atoms were constrained using the LINCS algorithm(41) to allow the time step of 2 fs. The desired temperature at either 310 K or 323 K was controlled with Nose-hoover thermostat(42, 43) (1.0 ps coupling constant). The pressure was maintained at 1 atm using the Parrinello-Rahman(44) semi-isotropic barostat with a coupling constant of 1.0 ps. Equilibrium MD systems were performed with two repeats. Each repeat was run with different initial velocities. The summary of all production runs is shown in Suppl. Table 2. The results were analysed using built-in function in GROMACS package. Molecular graphic pictures were prepared using VMD.

## Acknowledgments

We are thankful for financial support through BBSRC project BB/R019061/1. We acknowledge Diamond Light Source for access and support of the cryo-EM facilities at the UK’s national Electron Bio-imaging Centre (eBIC) [under proposal EM-19832]. We thank Emma Hasketh and Rebecca Thompson from the Astbury Centre for Structural Molecular Biology Leeds for support during measurements. We acknowledge support from LonCEM, King’s College London during measurements. The University of Sheffield FoS cryo-EM facility was used for grid preparation and optimization. We are grateful to Justin Kollman for help with the initial stages of this project.

## Notes

### Competing Interest Statement

The authors have declared no competing interest.

