## Supplementary material for "Structure and lipid dynamics in the *A. baumannii* maintenance of lipid asymmetry (MLA) inner membrane complex": Suppl. Fig.

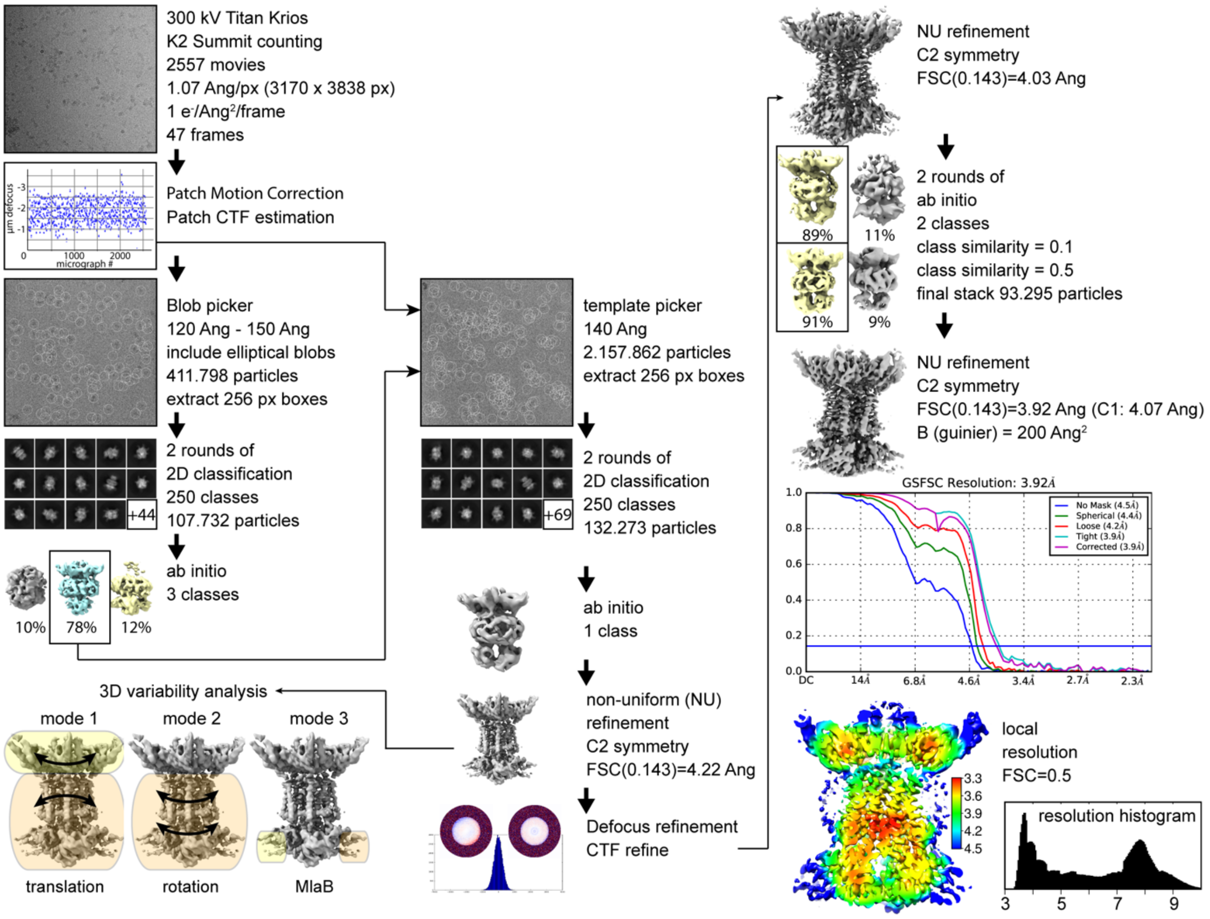


**Supplemental Figure 1:** processing of the MlaBDEF-AppNHp dataset in CryoSPARC v2.14.2. 2557 images were recorded at 300 kV on a Titan Krios with an energy filtered Gatan BioQuantum967 detector (K2 summit) with a Nyquist frequency of 2.14 Ang. Patch motion correction resulted in a detected defocus range of -1 to -2.5 µm. Initial blob picking with a diameter range from 120-150 Ang including elliptical shapes were extracted and 2D classified two times into each 250 classes. Ab initio 3D models were generated and the best class was used to re-pick particles template-based. After two rounds of 2D classification an ab initio 3D model was generated and refined using CryoSPARCs Non-Uniform (NU) refinement procedure with C2 symmetry, followed by global and local CTF refinement and two rounds of ab initio model generation with class similarity values of 0.1 and 0.5, respectively. The final particle stack contained 93.295 particles and was NU refined to 3.92 Ang with C2 symmetry (4.07 Ang with C1 symmetry). The map was sharpened with the Guinier plot B-factor of -200 Ang$^2$. Fourier Shell Correlation plot is indicated as well as local resolutions at FSC=0.5 projected on the final map. A histogram with the full local resolution range is also indicated. The first high resolution 3D structure was used as an input for CryoSPARCs 3D variability jobtype with 6 modes. Only the first three modes showed global changes; firstly, translation of MlaD against MlaBEF (Supplemental Movie 1), secondly, rotation of the MlaBEF part against MlaD (Supplemental Movie 2) and thirdly, alternating appearance of MlaB, indicating lower occupancy of this part compared to the MlaDEF part.

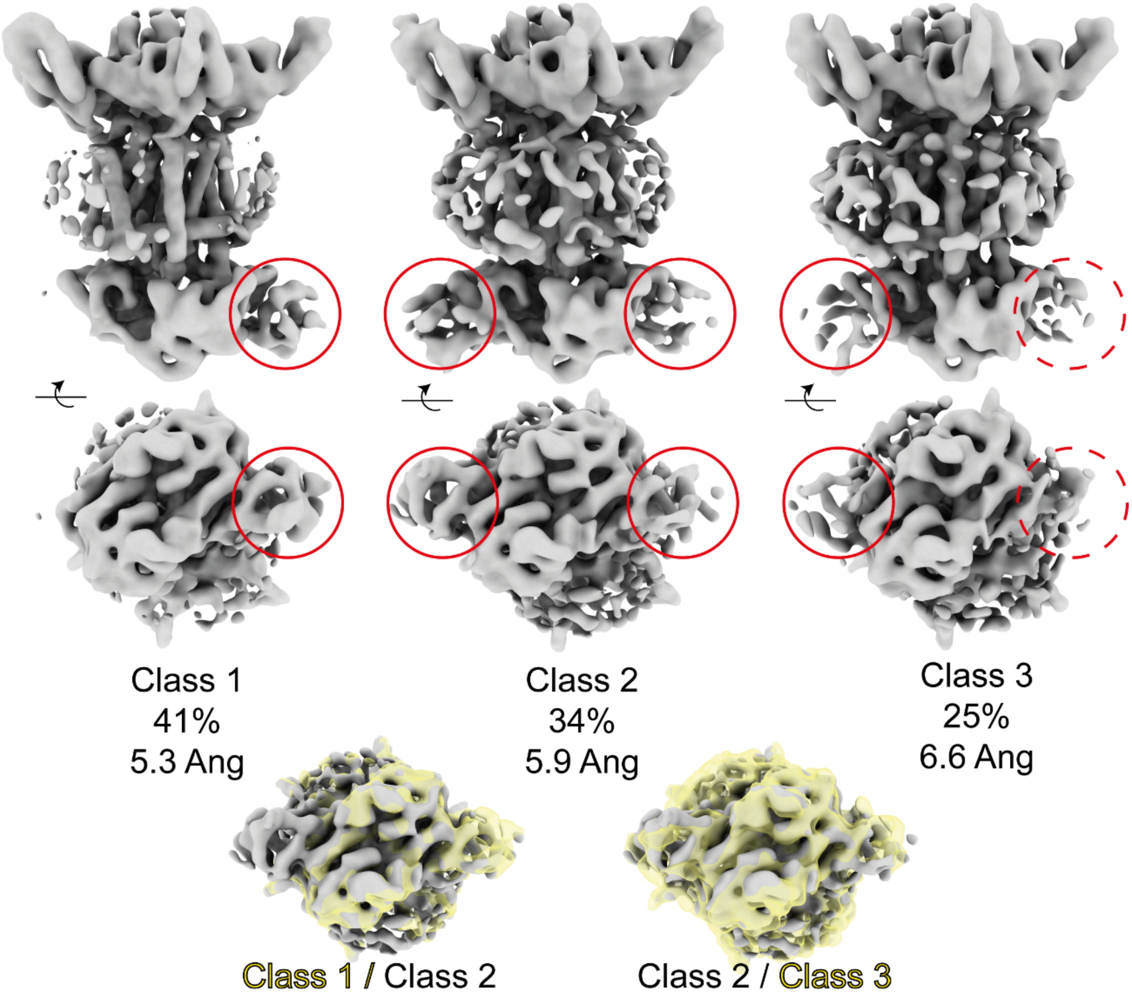


**Supplemental Figure 2:** MlaB binding (red circles) occurred on both binding sites in about 50% of the particles (classes 2 and 3) and on only one binding site in the other 50% of the particles (class 1). Maps were obtained by Non-Uniform refinement in C1 symmetry after heterogeneous refinement with 3 classes in CryoSPARC. Alignments of Classes 1-3 snow no major structural changes upon MlaB binding.

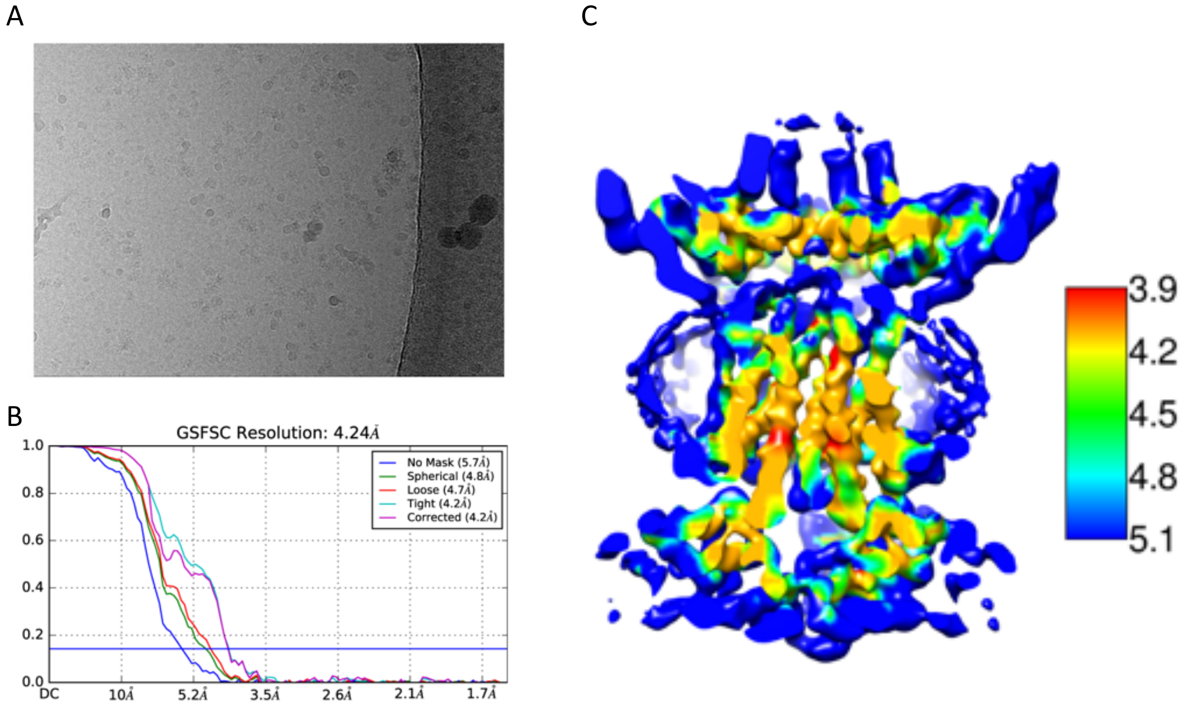


**Supplemental Figure 3:** processing of the apo MlaBDEF$_{ab}$ dataset in CryoSPARC v2.14.2. Images were recorded on 300 kV Titan Krios instruments equipped with Gatan K3 Bioquantum detector in counting mode.. After blob picking and 2D classification selected 2D classes were used for template picking. After several rounds of ab initio 3D structure generation and 3D classification, non-uniform refinement with C2 symmetry led to a map with a global resolution of ~ 4.2 Å.

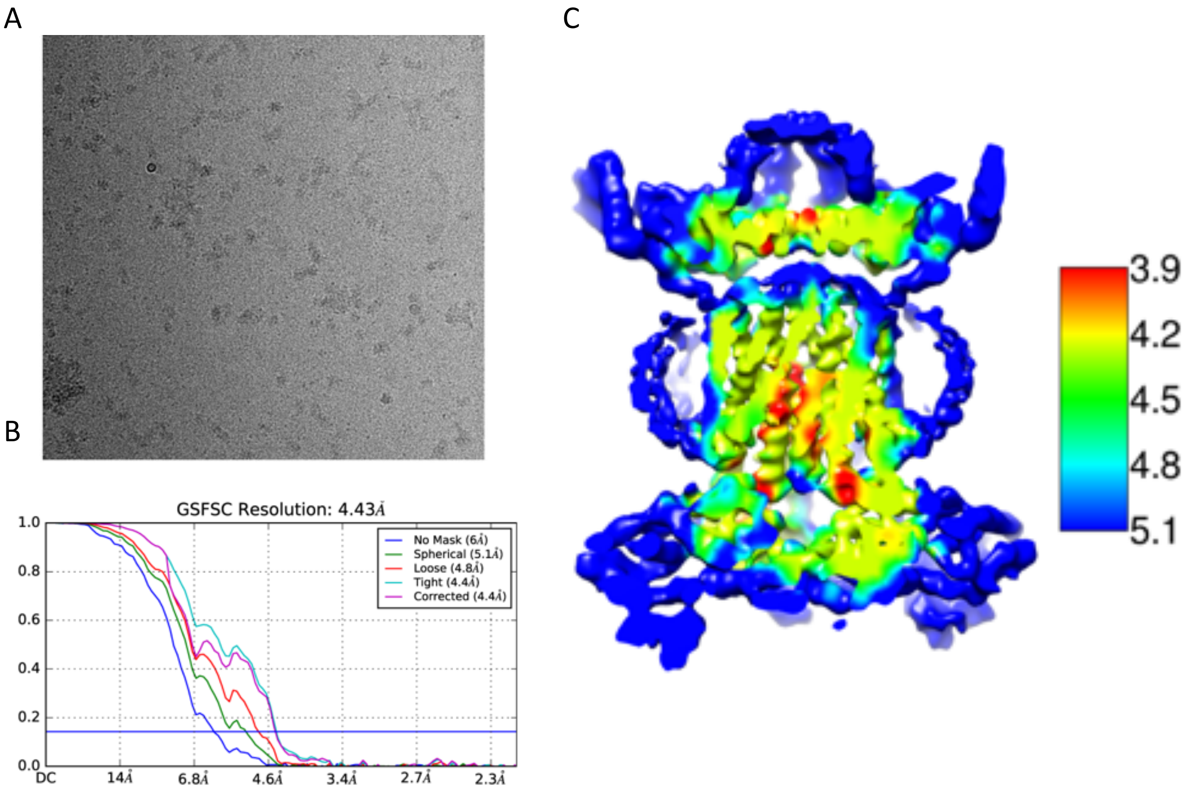


**Supplemental Figure 4:** processing of the MlaBDEF$_{ab}$ -ADP dataset in CryoSPARC v2.14.2. Images were recorded on 300 kV Titan Krios instruments equipped with Gatan K2 Summit detector in counting mode. After blob picking and 2D classification selected 2D classes were used for template picking. After several rounds of ab initio 3D structure generation and 3D classification, non-uniform refinement with C2 symmetry led to a map with a global resolution of ~ 4.4 Å.

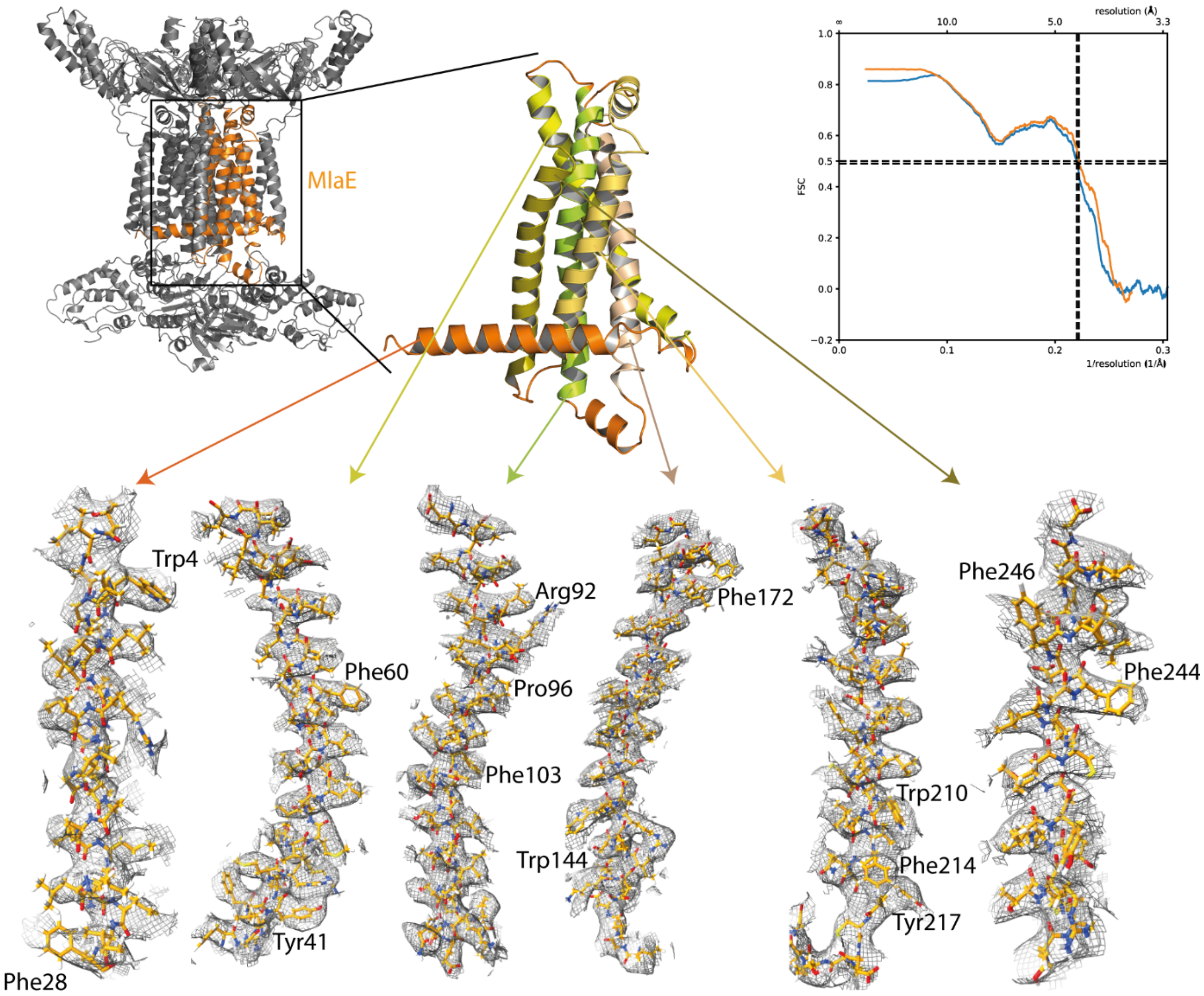


**Supplemental Figure 5:** de novo model building of MlaE (orange). Large side chains that allowed sequence mapping are indicated as well as map-to-model FSC of the whole $MlaBDEF_{ab}$ protein complex.

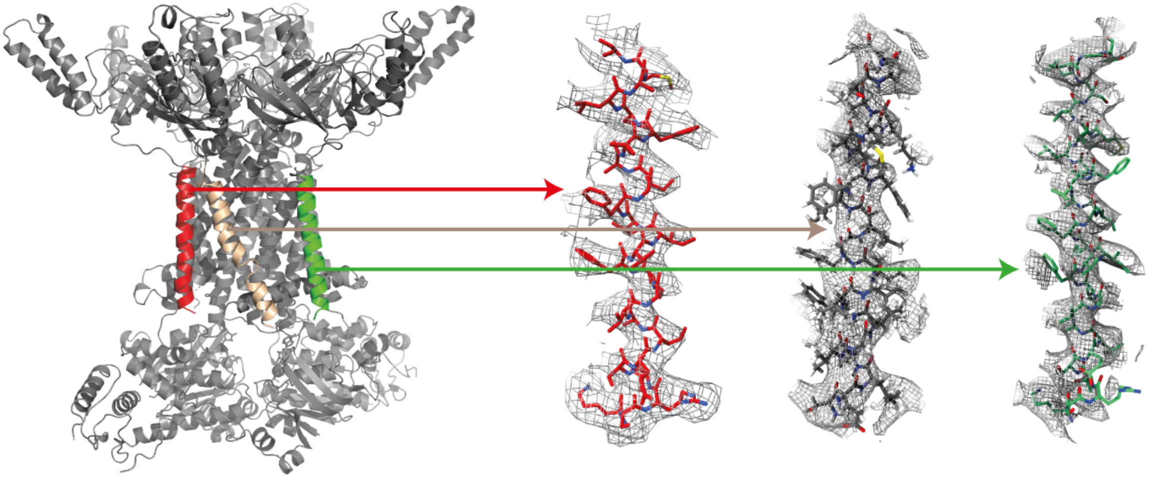

**Supplemental Figure 6:** Enclosed N-Helix of MlaD is significantly better resolved compared to peripheral MlaD N-helices (grey, green).

Supplementary Table 1 Equilibrium Protocol

| Step | Time Step (fs) | Total Time (ns) | Protein Backbone Restraints (kJ $mol^{-1}$ nm $^{-2}$) | Protein Sidechain Restraints (kJ $mol^{-1}$ nm $^{-2}$) | Ensemble |
|---|---|---|---|---|---|
| 1 | 1 | 0.125 | 4000 | 2000 | NVT |
| 2 | 1 | 0.125 | 2000 | 1000 | NVT |
| 3 | 1 | 0.125 | 1000 | 500 | NPT |
| 4 | 2 | 0.5 | 500 | 200 | NPT |
| 5 | 2 | 0.5 | 200 | 50 | NPT |
| 6 | 2 | 20 | 50 | 0 | NPT |

Supplementary Table 2: Summary of the equilibrium MD simulation systems

| System | Substrate | Temperature (K) | Simulation Length (ns) |
|---|---|---|---|
| *Apo* Mla | - | 310 | 500 ($\times$ 2) |
| *Apo* Mla | - | 323 | 500 ($\times$ 2) |
| *Holo* Mla | 7 POPE | 310 | 500 ($\times$ 2) |
